# Identification of a divergent botulinum neurotoxin like gene cluster in *Furfurilactobacillus*

**DOI:** 10.1101/2025.01.16.633459

**Authors:** Xin Wei, Pyung-Gang Lee, Min Dong, Andrew C. Doxey

## Abstract

Botulinum neurotoxins (BoNTs) are among the most potent biological toxins and are traditionally associated with *Clostridium* species. However, recent discoveries have identified BoNT-like proteins in diverse bacterial genera, revealing an expanding family of neurotoxins with unique evolutionary and functional traits. In this study, we report the identification of a novel BoNT-like gene cluster in *Furfurilactobacillus* sp. OKN36, encoding a toxin we tentatively designate as “Furfuritoxin”. Sequence and structural analyses indicate that Furfuritoxin shares key domains with clostridial neurotoxins, including a light chain zinc metalloprotease domain and heavy chain translocase and binding domains, while exhibiting significant sequence divergence. Phylogenetic analysis places Furfuritoxin within a divergent lineage alongside previously reported BoNT/Wo, suggesting a shared ancestral relationship. Gene neighborhood analysis reveals features shared with other BoNT gene clusters including ORFX-related genes and conjugation-associated elements, indicating horizontal gene transfer may have facilitated its distribution. This study adds to the growing family of BoNT-like toxins, providing insights into their evolution and diversity.

## Introduction

Clostridial neurotoxins (CNTs), including botulinum neurotoxin (BoNT) and tetanus neurotoxin (TeNT), are among the most powerful biological toxins identified to date. These neurotoxins are the primary agents responsible for the neuroparalytic conditions botulism and tetanus. Decades of extensive research have positioned these toxins as central models for understanding the molecular mechanisms of infectious diseases caused by toxin-producing organisms^1–5^.

Botulinum neurotoxins (BoNTs) and tetanus neurotoxins (TeNTs) act by disrupting neurotransmitter release, though their effects differ due to distinct cellular targeting. Both toxins are internalized into neurons via receptor-mediated endocytosis, where their light-chain protease domains are translocated into the cytosol. BoNTs cleave SNARE proteins, such as VAMP, syntaxin, and SNAP25, within motor neurons, preventing synaptic vesicle fusion and causing flaccid paralysis. In contrast, TeNT undergoes retrograde transport to the soma of motor neurons in the spinal cord, where it blocks inhibitory neurotransmitter release, resulting in spastic paralysis. Both mechanisms lead to fatal respiratory failure if untreated.

Structurally, BoNT and TeNT are organized into three functional domains^5^. The light chain (LC) harbors the protease activity that cleaves SNAREs. The heavy chain (HC) is split into two regions: the translocation domain (HN), responsible for delivering the LC into the cytosol, and the binding domain (HC), which facilitates attachment to various host receptors in a serotype-specific manner. Historically, BoNTs have been grouped into seven serotypes (A–G)^6–9^. Serotypes A, B, and E are linked to human botulism, while serotypes C, D, and the hybrid CD primarily affect wildlife. The host range for BoNT/G remains poorly defined. BoNTs are encoded by different mechanisms, including chromosomal genes, plasmids, or phages, depending on the subtype, and non-toxic strains lacking BoNTs are common. These toxins are not exclusive to *C. botulinum*, being also produced by *C. noyvi, C. baratii*, and *C. butyricum*, among others.

With the rapid growth of publicly available genomic data, researchers have discovered a growing number of BoNT-like toxins through computational analyses^8,10–18^. A landmark discovery was that of Mansfield et al.^11^, who identified the first BoNT homolog outside the genus *Clostridium*, in *Weissella oryzae*. This toxin, called “BoNT/Wo”, has been reported to cleave VAMP at an unconventional site^19^, though its specificity remains to be established. Structural analyses revealed that BoNT/Wo retains the domain organization characteristic of BoNTs^11^, but phylogenetic analysis places it on a branch outside the canonical BoNT clade, implying that the canonical BoNTs (A–G) represent a subgroup of a larger family of BoNT-like proteins. Later, Zhang et al.^15^ described BoNT/X, a unique member of the BoNT family that cleaves multiple VAMP isoforms and several non-traditional substrates. Subsequently, in 2018, a BoNT homolog, termed BoNT/En, was identified in the commensal and opportunistic pathogen *Enterococcus faecium*^13^. BoNT/En cleaves both SNAP25 and VAMP2 at distinct sites and is carried on a conjugative plasmid, suggesting horizontal gene transfer among *Enterococcus* strains. Phylogenetically, BoNT/En groups closely with BoNT/X and shares genomic features. Following the identification of BoNT/En, Contreras et al. identified a BoNT-like toxin in *Paraclostridium bifermentans*^14^. Contreras et al. demonstrated that PMP1 acts as an insecticidal toxin, cleaving syntaxin 1 in mosquitoes. PMP1 clusters within the BoNT/X lineage alongside BoNT/En. Genomic analyses revealed the presence of adjacent ORFX genes, which may enhance PMP1’s toxicity by facilitating absorption in the insect gut. In 2022, Wei et al.^16^ then identified a unique BoNT-like gene cluster in two strains of *Paeniclostridium ghonii*. The identified PG-toxins are unique by splitting their heavy and light chains into separate genes, and have a gene neighborhood structure suggesting an insecticidal function^16^. Finally, the most recent addition to the BoNT family tree was identified in 2023 in *Bacillus toyonensis*^17^. Similar to that of PG toxin (PGT), this toxin (BTNT) was found to cluster phylogenetically with the X/En/PMP1 lineage.

Multiple lines of evidence suggest that the X/En/PMP1/BTNT/PGT clade of BoNT-like toxins may be a sister lineage to the canonical BoNTs which have evolved a unique specificity for invertebrate hosts such as insects^8,14^. Although studies are shedding light on the biology of this lineage of BoNTs, very little is known about the highly divergent BoNT/Wo toxin, which falls outside of this group. Here, we report the identification of a similarly divergent toxin to BoNT/Wo, in the organism, *Furfurilactobacillus* sp. OKN36. We analyze this new BoNT-like toxin gene cluster in terms of its sequence and phylogenetic relationships to other BoNTs, and gene neighborhood. This work adds to the growing diversity of BoNT-like toxins, which may lead to new functional insights and reveal novel host specificities.

## Results and Discussion

### Identification of a BoNT-related sequence in Furfurilactobacillus sp. OKN36

On Nov 24, 2024, we performed a BLAST-based search for BoNT-like toxins in the NCBI database. Our search, using BoNT/Wo (accession # WP_027699549.1) as a query identified a previously unidentified match (WP_407880486.1) in the genome of the bacterium, *Furfurilactobacillus* sp. OKN36. This sequence was the closest match to BoNT/Wo, with 28.6% sequence identity and an *E*-value of 3 × 10^−168^. Similarly, a reciprocal search done using the *Furfurilactobacillus* sequence (WP_407880486.1) as a query identified BoNT/Wo as the closest match, confirming a relationship between these two toxins.

The *Furfurilactobacillus* sequence (1347 aa in length) is annotated in the NCBI as a “LamG-like jellyroll fold domain-containing protein” and, based on NCBI Conserved Domain Database annotation, contains an N-terminal Peptidase_M27 domain family and C-terminal Laminin_G_3 domain homologous to similar domains in clostridial neurotoxins. The N-terminal zinc protease domain contains an <u>HE</u>LI<u>H</u> motif, suggesting that it is an active metalloprotease. The AlphaFold3^20^ structure revealed an overall fold and domain architecture characteristic of clostridial neurotoxins, with some regions (i.e., the central translocation domain) resolving more poorly in the model (**Figure 1**). We therefore designated this protein as “Furfuritoxin”.

**Figure 1.**
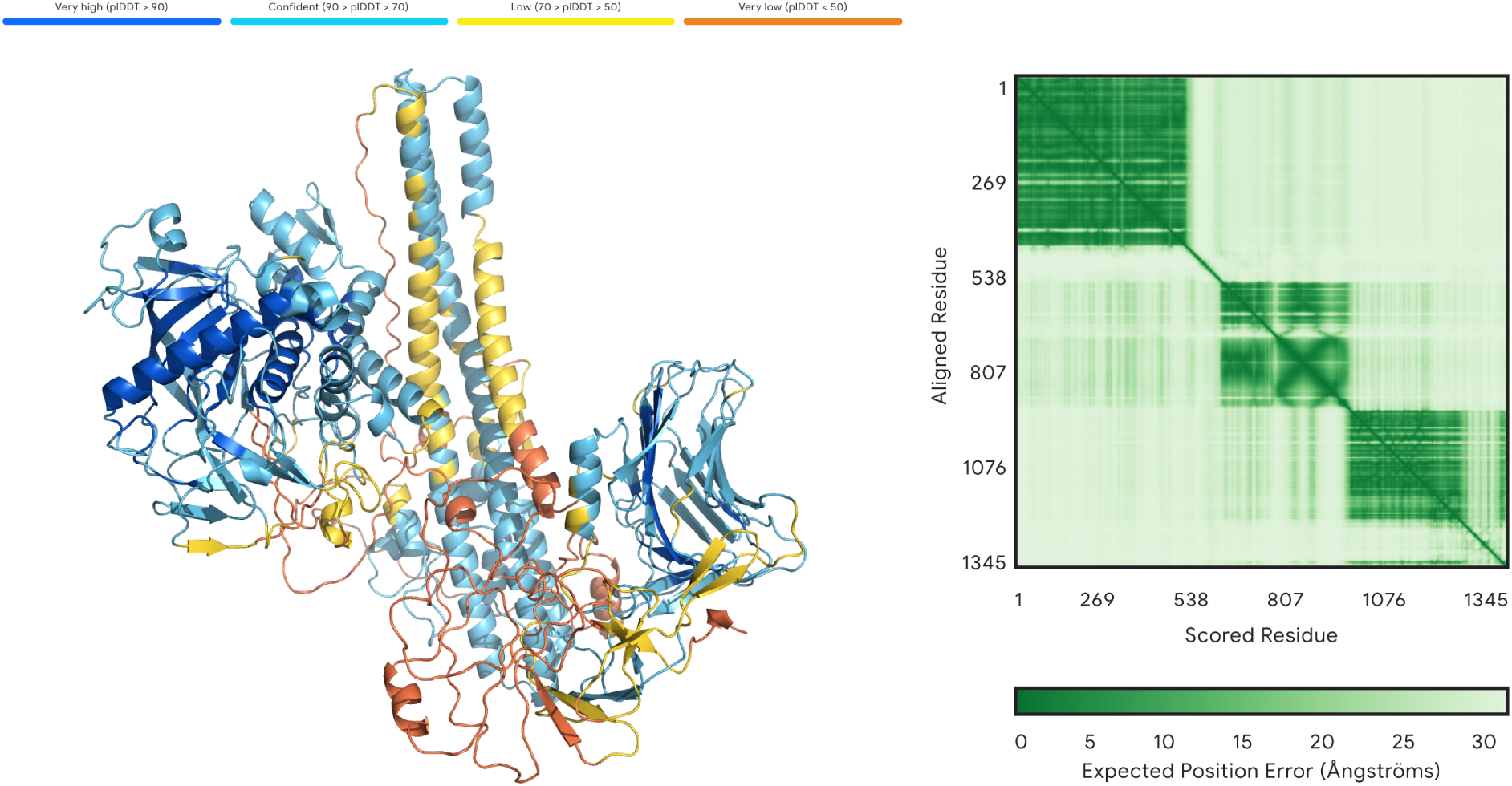
Predicted structure of a BoNT-like toxin from *Furfurilactobacillus* sp. OKN36, using AlphaFold3.

### Gene neighborhood analysis

The gene neighborhood of the newly identified toxin gene in *Furfurilactobacillus* is shown in **Figure 2** along with other BoNT gene clusters. The genomic context of the *Furfurilactobacillus* BoNT-like gene exhibits numerous similarities to other BoNT gene clusters. BLAST analysis of the upstream region of the new toxin revealed P47/ORFX-containing proteins with homology to ORFX2, ORFX3, and P47 proteins encoded in other BoNT gene neighborhoods. Notably, a potential duplication of the ORFX2 gene was identified in the upstream region of the new toxin. The rest of the neighboring gene content aligns more closely with that in the *Weissella oryzae* gene cluster. Conjugation-related genes, including *traG* (conjugal protein) and *mobP2* (relaxase), were found in *Furfurilactobacillus* and *Weissella oryzae*, respectively. Additionally, *topA* (topoisomerase I) was identified in both *Furfurilactobacillus* and *Weissella oryzae*.

**Figure 2.**
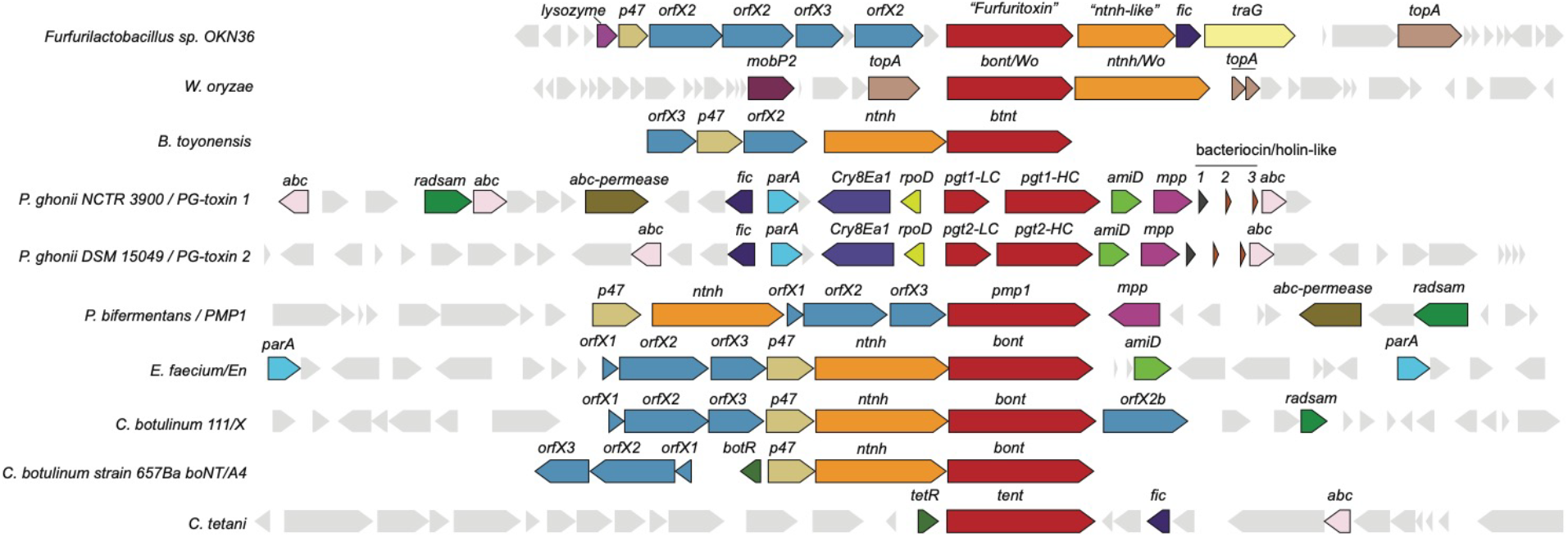
Gene neighborhood structure of a BoNT-like gene cluster in *Furfurilactobacillus* sp. OKN36 and relationship to other BoNT gene clusters.

The presence of the *traG* gene in the gene neighborhood of the new toxin suggests that the DNA segment containing the toxin may have been transferred from another CNT-containing bacterium through conjugation. Also, a lysozyme-associated protein in the upstream region may play a role in bacterial cell wall degradation, potentially facilitating toxin release or horizontal gene transfer. The topoisomerase *topA* gene in the downstream region that modifies the topology of DNA could contribute to DNA recombination events. Together, these neighboring genes suggest that the new toxin in *Furfurilactobacillus* may have been acquired from a different CNT-containing bacterium, potentially through horizontal gene transfer mechanism.

### Phylogenetic analysis

To investigate the evolutionary relationships between the new toxin and other BoNTs, as well as previously identified BoNT-like toxins (PG toxins, BTNT, Wo, etc.), a maximum-likelihood phylogenetic tree was constructed using IQ-TREE (**Figure 3**). The analysis revealed that the new toxin forms a distinct clade together with BoNT/Wo, supported by strong (100%) bootstrap values. This finding suggests that the new toxin and BoNT/Wo share a common ancestor. However, the considerable sequence divergence and long evolutionary branch connecting these toxins indicates a potentially ancient relationship or, alternatively, strong selective pressures driving accelerated rates of mutation.

**Figure 3.**
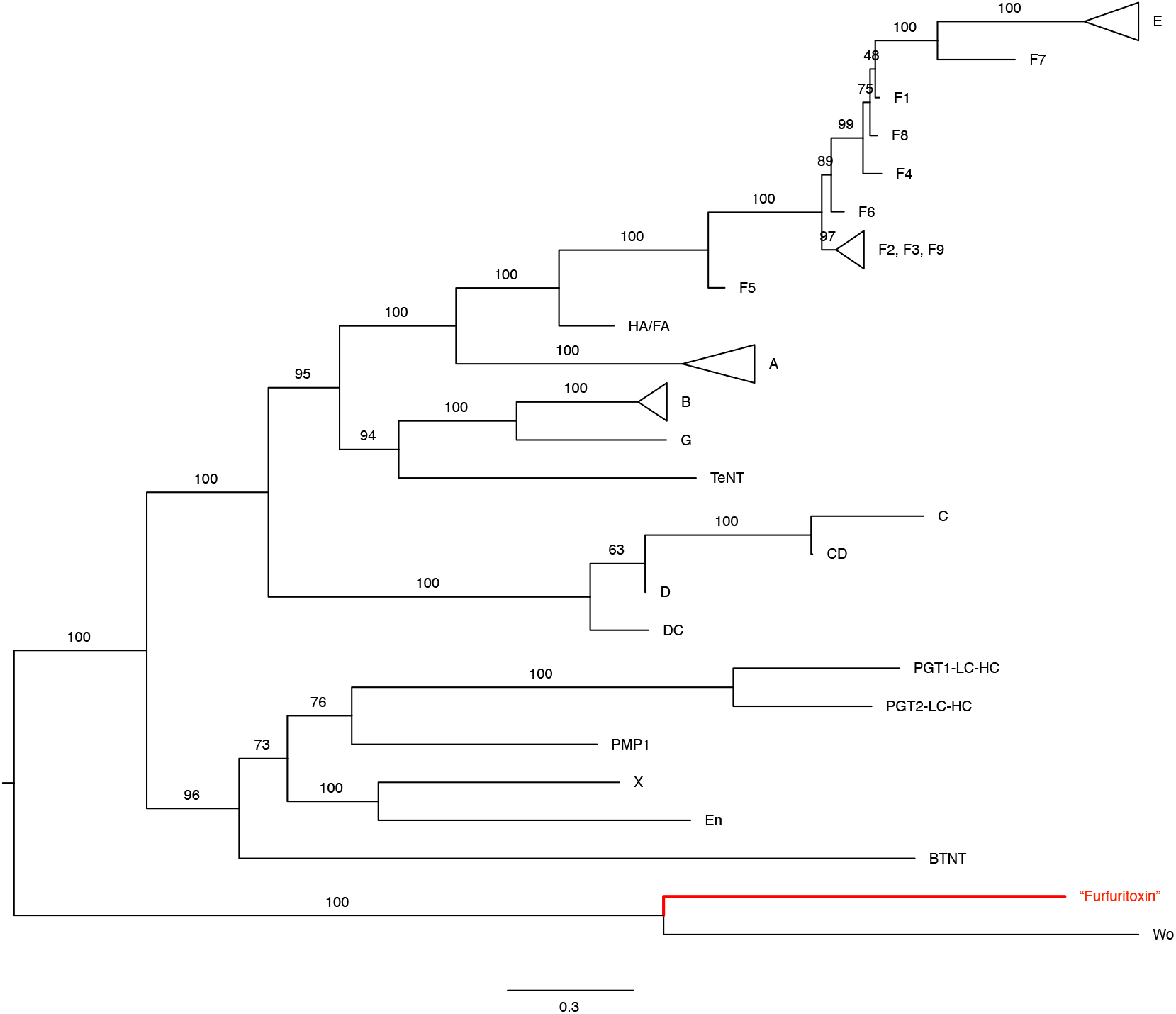
Maximum-likelihood phylogenetic tree of the BoNT family including the newly identified*Furfurilactobacillus* sequence (red).

## Methods

### Gene neighborhood analysis

Gene neighborhood visualization was done using the AnnoView^21^ web server at annoview.uwaterloo.ca. Specifically, GBK files of genomic regions containing the BoNT genes were retrieved from the NCBI and uploaded to AnnoView. Genes with unknown functions were annotated manually by performing BLAST searches against the NCBI non-redundant protein database.

### Phylogenetic analysis

A curated list of known BoNT protein sequences was compiled from bontbase.org and included previously discovered toxin genes (PG toxins and BTNT). PGT1-LC-HC and PGT2-LC-HC, were generated by concatenating PGT1-LC with PGT1-HC and PGT2-LC with PGT2-HC, respectively. Multiple sequence alignment of all protein sequences was performed using MAFFT v7.505^22^ with the L-INS-i algorithm, which is optimized for sequences with global homology. A phylogenetic tree of all BoNT toxins was constructed using IQ-TREE version 2.3.6^23^. We then midpoint rooted the tree and collapsed clades that include toxins from the same subtype.

